# Small, correlated changes in synaptic connectivity may facilitate rapid motor learning

**DOI:** 10.1101/2021.10.01.462728

**Authors:** Barbara Feulner, Matthew G. Perich, Raeed H. Chowdhury, Lee E. Miller, Juan Álvaro Gallego, Claudia Clopath

## Abstract

Animals can rapidly adapt their movements to external perturbations. This adaptation is paralleled by changes in single neuron activity in the motor cortices. Behavioural and neural recording studies suggest that when animals learn to counteract a visuomotor perturbation, these changes originate from altered inputs to the motor cortices rather than from changes in local connectivity, as neural covariance is largely preserved during adaptation. Since measuring synaptic changes *in vivo* remains very challenging, we used a modular recurrent network model to compare the expected neural activity changes following learning through altered inputs (H_input_) and learning through local connectivity changes (H_local_). Learning under H_input_ produced small changes in neural activity and largely preserved the neural covariance, in good agreement with neural recordings in monkeys. Surprisingly given the presumed dependence of stable neural covariance on preserved circuit connectivity, H_local_ led to only slightly larger changes in neural activity and covariance compared to H_input_. This similarity is due to H_local_ only requiring small, correlated connectivity changes to counteract the perturbation, which provided the network with significant robustness against simulated synaptic noise. Simulations of tasks that impose increasingly larger behavioural changes revealed a growing difference between H_input_ and H_local_, which could be exploited when designing future experiments.

## Introduction

Animals, particularly primates, can perform a great variety of behaviours, which they are able to adapt rapidly in the face of changing conditions. Since behavioural adaptation can happen even after a single failed attempt [Thoroughman and Shadmehr, 2000], the neural populations driving this process must be able to adapt equally fast. How this occurs remains unexplained [Sohn et al., 2020]. Rapid motor learning is typically studied using external perturbations such as a visuomotor rotation (VR), which rotates the visual feedback about the movement. Both humans and monkeys can learn to compensate for the error between actual and expected visual feedback in a few tens of trials [Wise et al., 1998, Krakauer et al., 2000]. Behavioural adaptation is accompanied by changes in the activity of neurons in primary motor cortex (M1) [Paz et al., 2003], and the upstream dorsal premotor cortex (PMd) [Wise et al., 1998]. It is unclear whether these neural activity changes are mediated by synaptic weight changes within the motor cortices or are driven by altered inputs from even further upstream areas.

When learning a skill over many days, behavioural improvements are paralleled by rewiring between M1 neurons [Rioult-Pedotti et al., 1998, Kleim et al., 2004, Xu et al., 2009, Roth et al., 2020]. This seems not to be the case for rapid learning: throughout VR adaptation, the statistical interactions across neural populations in both M1 and PMd remain preserved [Perich et al., 2018]. These preserved interactions rule out any large synaptic changes within the motor cortices, as they would cause these models to degrade [Gerhard et al., 2013, Rebesco et al., 2010]. Instead, rapid VR adaptation may be driven by the cerebellum [Tseng et al., 2007, Rabe et al., 2009, Schlerf et al., 2012, Tzvi et al., 2020] and/or posterior parietal cortex [Diedrichsen et al., 2005, Tanaka et al., 2009].

A pioneering Brain Computer Interface (BCI) study cast further doubt that significant synaptic changes occurring within M1 are necessary for rapid learning [Sadtler et al., 2014, Golub et al., 2018]. In that study, monkeys controlled a computer cursor linked by a “decoder” to the activity of recorded M1 neurons. After learning to use a decoder that directly mapped ongoing neural activity onto cursor movements, the monkeys were exposed to one of two types of perturbations. When faced with a new decoder that preserved the statistical interactions (i.e., neural covariance) across M1 neurons, the monkeys could master it within minutes. In stark contrast, if the new decoder required changes in the neural covariance, they could not learn it within one session – in fact, it required a progressive training procedure spanning just over nine days on average [Oby et al., 2019].

Recording synaptic changes on a large scale in-vivo remains extremely challenging and has not been achieved during rapid motor learning. Recurrent neural network (RNN) models offer an exciting yet unexplored opportunity to test the effect of synaptic changes - weight changes in the model - on simulated activity during motor learning. RNNs trained on motor, cognitive and BCI tasks exhibit many striking similarities with the activity of neural populations recorded in animal studies [Mante et al., 2013, Sussillo et al., 2015, Rajan et al., 2016, Song et al., 2017, Wang et al., 2018, Michaels et al., 2020, Perich et al., 2021], suggesting a fundamental similarity between the two. Previous work using RNNs to model the BCI experiment described above [Sadtler et al., 2014] showed that network covariance can be highly preserved even when learning is happening through weight changes within the network [Feulner and Clopath, 2021]. Thus, contrary to widespread intuition, functionally relevant synaptic weight changes may not necessarily lead to measurable changes in statistical interactions across neurons [Das and Fiete, 2020]. Therefore, there might be synaptic changes within PMd and M1 during VR adaptation that are very hard to identify through the analysis of neural population recordings.

Here, we used RNN models to test whether VR adaptation might be mediated by synaptic changes within PMd and M1 that largely preserve the neural covariance within these areas. We addressed this question by comparing how adaptation based on connection weight changes (H_local_) within PMd and M1 alters network activity compared to the corresponding activity changes if VR adaptation is based on altered inputs from upstream areas (H_input_) [Tseng et al., 2007, Rabe et al., 2009, Schlerf et al., 2012, Tzvi et al., 2020, Diedrichsen et al., 2005, Tanaka et al., 2009, Perich et al., 2018] (Figure 1A). To validate our modelling results, we compared our simulations to experimental recordings from PMd and M1 populations during the same VR task [Perich et al., 2018].

**Figure 1:**
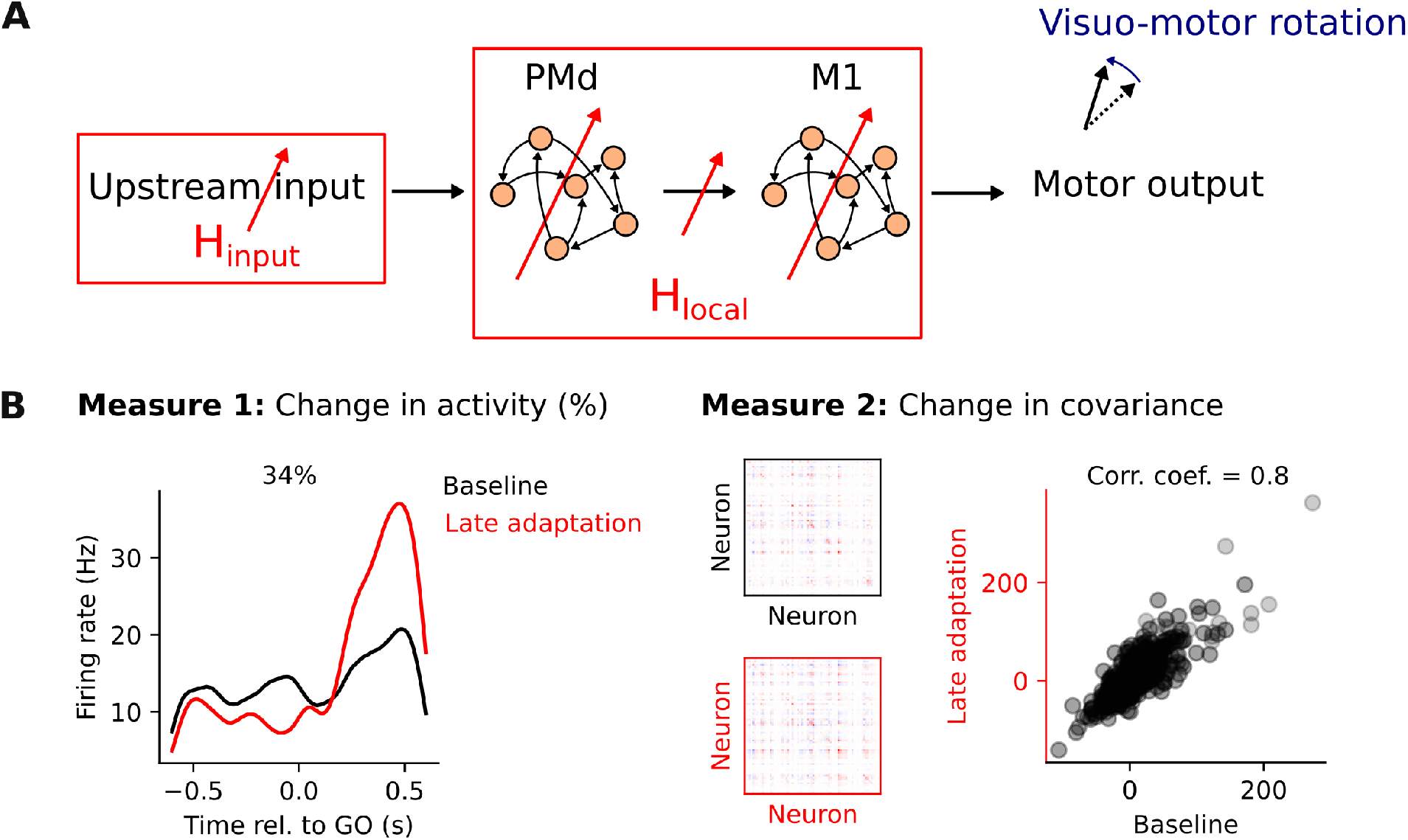
Competing hypotheses to explain where learning happens during a visuomotor rotation task. **A**. To study the processes mediating motor cortical activity changes during adaptation in a visuomotor rotation task, we analyse and model the activity of neural populations within dorsal premotor cortex (PMd) and primary motor cortex (M1). We compare two hypotheses: plasticity upstream of PMd/M1 (H_input_) and plasticity within PMd/M1 (H_local_). **B**. Measures to quantify the changes in neural activity following adaptation: 1) relative change in trial-averaged single neuron activity; 2) change in neural covariance. Both measures compare baseline trials to late adaptation trials, after monkeys had adapted to the task.

Under H_local_, the changes in network activity and covariance following VR adaptation only slightly exceeded those under H_input_ and were comparable to experimental observations. Thus, when using neural population recordings alone, it may be more challenging to disentangle these two hypotheses than previously thought. For both H_input_ and H_local_, the learned connectivity changes were small and highly coordinated, which made them surprisingly robust to noise. To identify additional differences between H_input_ and H_local_, we examined learning during paradigms requiring larger behavioural changes. Covariance changes were larger for these paradigms in both PMd and M1 under H_input_, but only in M1 under H_local_, thus providing a way to distinguish between the two hypotheses in future experiments. Our findings have implications for the interpretation of experimental neural activity changes during learning and suggest that tasks eliciting large behavioural changes may be necessary to elucidate how neural populations adapt their activity during rapid learning.

## Results

To understand whether motor adaptation could be driven by synaptic changes within PMd and M1, we simulated a VR adaptation task using a modular RNN that modelled these two areas and compared the resulting changes in network activity to those of neural population recordings from PMd and M1 during the same VR task [Perich et al., 2018]. We quantified neural activity changes both in the experimental data and in the model using two measures (Figure 1B): 1) the relative change in trial-averaged single neuron activity, and 2) the change in neural covariance (Methods). Combined, they capture aspects of single neuron as well as population-wide activity changes during adaptation.

### Small but measurable changes in neural activity within PMd and M1 during VR adaptation

Monkeys were trained to perform an instructed delay task, in which they reached to one of eight visual targets using a planar manipulandum to receive a reward (Methods). After performing a block of unperturbed reaches (200-243 trials, depending on the session), visual feedback about the position of the hand was rotated by 30° clockwise or counter-clockwise, depending on the session. Monkeys adapted rapidly to these perturbations: the curved reaches observed immediately after the perturbation onset became straighter after tens of trials, with the hand trajectories in the second (late) half of the adaptation block becoming more similar to the baseline trajectories (Figure 2A). The angular error quantifying the difference between initial reach direction and target location decreased during adaptation (Figure 2B). This error curve followed a similar trend for clockwise and counterclockwise perturbations, allowing us to analyse the different perturbations together.

**Figure 2:**
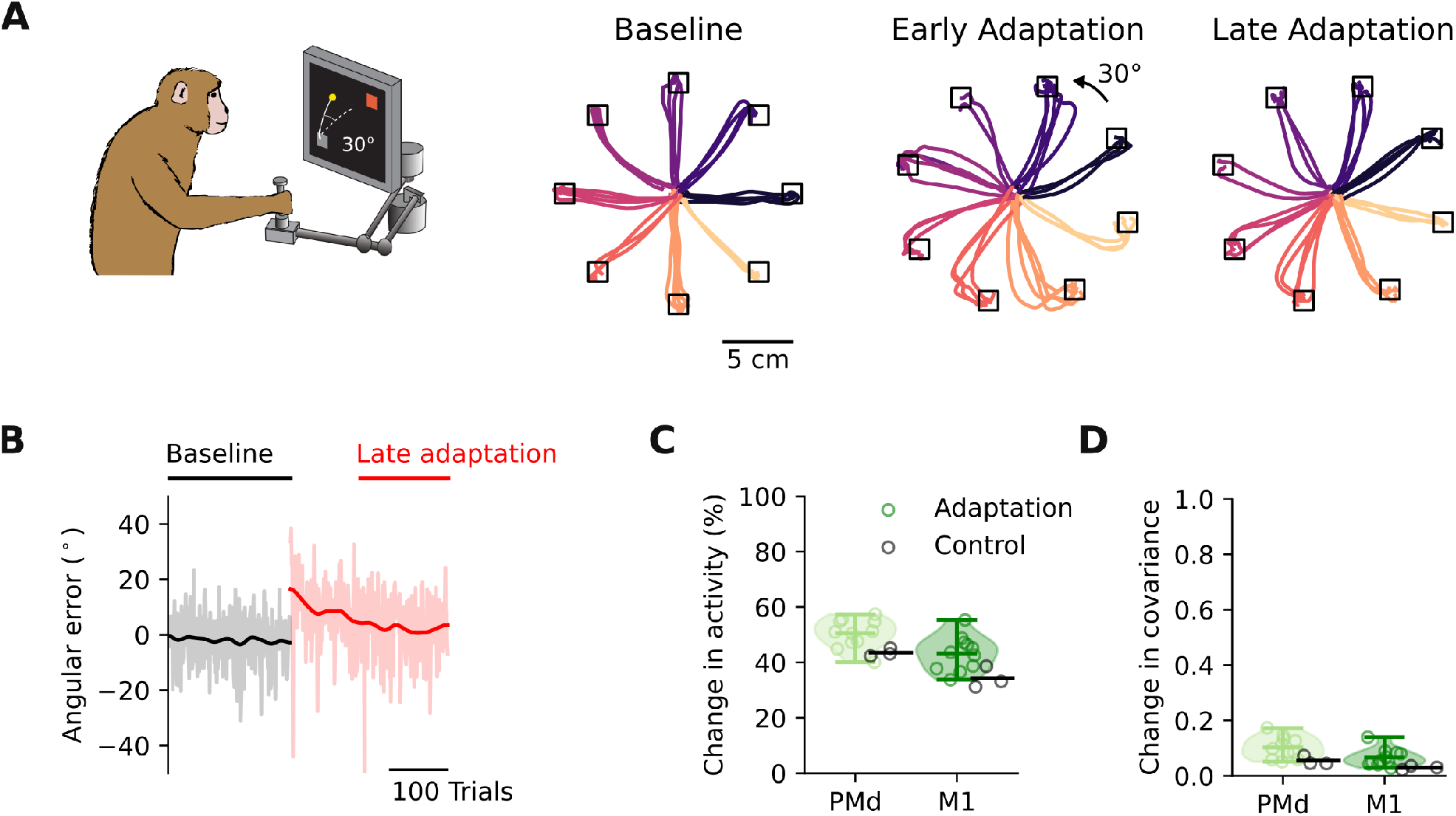
Small but measurable changes in neural activity within PMd and M1 during VR adaptation. **A**. Hand trajectories during the first 30 trials of the baseline, early adaptation (first 150 adaptation trials) and late adaptation epoch (last 150 adaptation trials). Trajectories are color-coded by target. Data from Monkey C. **B**. Angular error of the hand trajectories for the example session in (A) has the typical time course of adaptation. **C**. Change in trial-averaged activity following adaptation. Data pooled across all sessions from the two monkeys for PMd and M1 separately (green markers, 11 sessions). Control sessions during which no perturbation was applied are shown for comparison (black markers, 3 sessions). Shaded areas show estimate of data distribution and horizontal bars indicate mean and extrema. **D**. Change in covariance following adaptation. Same format as C.

Behavioural adaptation was accompanied by changes in neural activity within both PMd and M1 (Figure 2C) [Perich et al., 2018]. These changes exceeded those during control sessions, where no perturbation was applied (Figure 2C black; linear mixed model analysis: t=4.4, P=0.0017). The amount of change was greater within PMd than M1 (t=8.9, P*<*0.0001). We also found small but detectable changes in neural covariance during VR perturbation, suggesting that the statistical interactions across neurons change slightly during adaptation (Figure 2D). Again, these changes exceeded those of the control sessions (Figure 2D black; t=2.6, P=0.026).

### A modular recurrent neural network model to study VR adaptation

To test whether experimentally observed changes in motor cortical activity could be driven by rapid synaptic plasticity [Roth et al., 2020] within PMd and M1, we trained a modular RNN model [Sussillo et al., 2015, Michaels et al., 2020] to perform the centre-out reaching task that we studied experimentally. To mimic broadly the hierarchical architecture of the motor cortical pathways, input signals were sent to the PMd module which then projected to the M1 module, which produced the final output signal (Figure 3A; Methods). After initial training on the task, the model produced correct reaching trajectories to each of the eight different targets (Figure 3B), which had the same dynamics as the monkeys’ (Figure 3C). Furthermore, Principal Component Analysis (PCA; Methods) revealed that the population activity of the PMd and M1 network modules was similar to those of the corresponding recorded neural populations (Figure 3D-E). We used Canonical Correlation Analysis (CCA) to quantify this apparent similarity between model and experimental population activity (Methods): correlations were larger (more similar) when we compared the corresponding modelled and recorded brain areas than when comparing the two experimental brain areas with each other (Figure S1A). This indicates that the PMd and M1 modules are, respectively, more similar to their experimental counterparts than they are to each other, suggesting that our model captures well the main features of the actual data.

**Figure 3:**
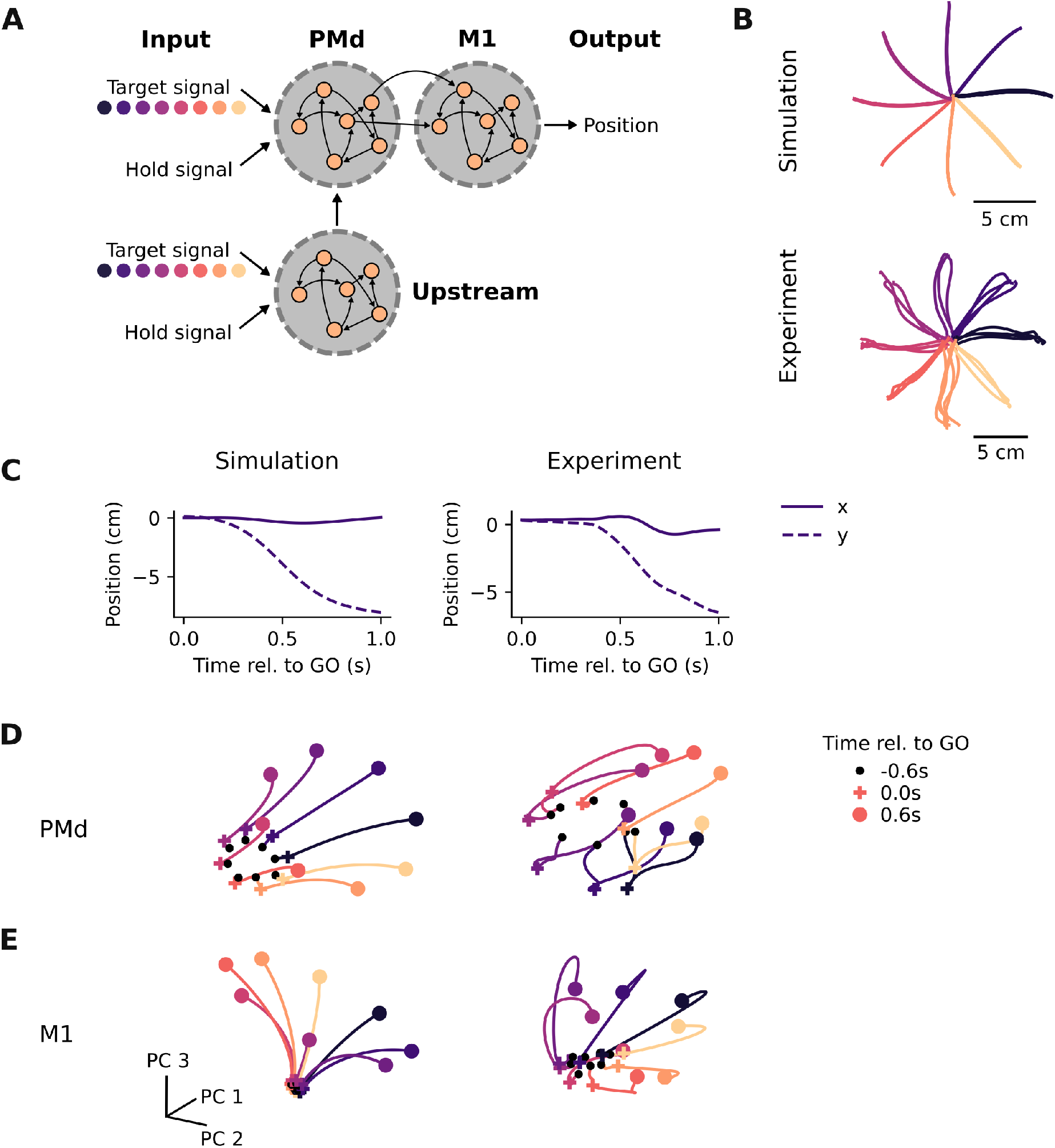
A modular recurrent neural network model to study VR adaptation. **A**. A modular RNN that models key motor cortical areas to study adaptation. **B**. Simulated (top) and actual (bottom) hand trajectories during 30 reaches to each target taken from one session from Monkey C. **C**. Example simulated and actual hand trajectories to one target. Note the similarity in kinematics between the model and the experimental data. **D**. Simulated PMd population activity recapitulates key features of actual PMd population activity. Neural trajectories go from 600 ms before the go cue (black dots) to 600 ms after the go cue (coloured dots); go cue is indicated with coloured crosses. Reaching targets are colour-coded as in B. **E**. Same as D for M1.

### Motor adaptation through altered inputs matches neural recordings

After having verified that our modular RNN recapitulates the key aspects of PMd and M1 population activity during reaching, we simulated the VR adaptation experiment. The model was retrained to produce trajectories rotated by 30°, replicating the perturbation monkeys had to counteract. Having full control of where learning occurred, we first constrained it to happen upstream of PMd (H_input_). As anticipated from previous modelling [Tanaka 2009] and experimental work [Perich et al., 2018], changes in areas upstream of the motor cortices can lead to successful adaptation: the hand trajectories produced after learning were correctly rotated by 30° to counteract the perturbation (Figure 4A).

**Figure 4:**
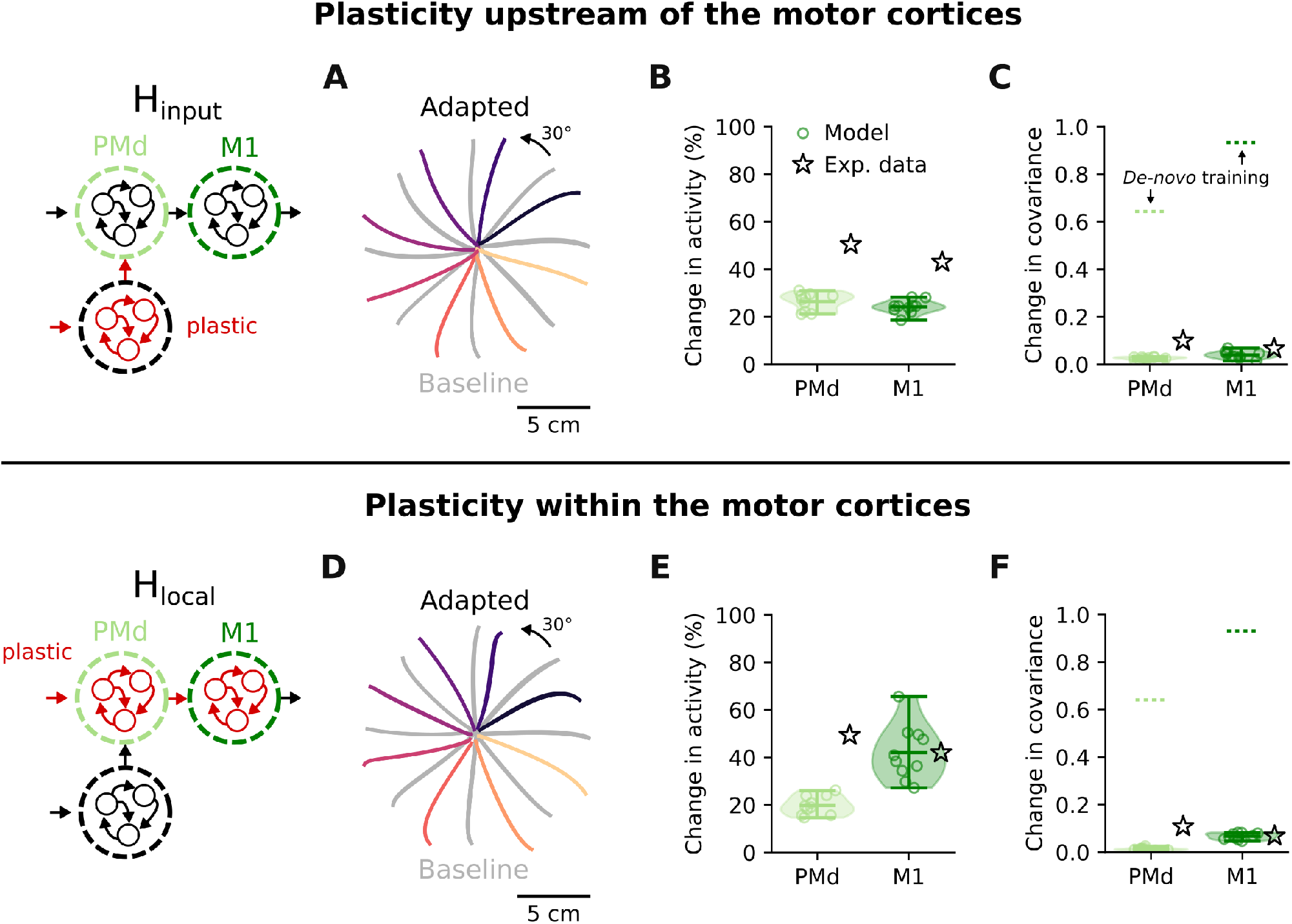
Activity changes following learning upstream (H_input_) and within (H_local_) the motor cortices. **A**. Hand trajectories after learning under H_input_ (coloured traces; baseline trajectories are shown in grey). **B**. Changes in trial-averaged activity following adaptation under H_input_ (green markers) for PMd and M1, and reference mean experimental values (black stars). **C**. Change in covariance following adaptation under H_input_, and reference values for change in covariance following the initial *de-novo* training (dashed lines). Data presented as in B. **D**. Hand trajectories after learning under H_local_. **E**. Change in trial-averaged activity following adaptation under H_local_. **F**. Change in covariance following adaptation under H_local_. Data in D,E,F are presented as in A,B,C.

When examining the activity of each of the PMd and M1 modules, the relative change in network activity was similar in magnitude to the changes observed in the corresponding neural population recordings (Figure 4B). PMd activity changed slightly more than M1 activity (Figure 4B), indicating a gradient between the two modules that was also present in the experimental data (Figure 2C). When looking at the change in interactions between neurons, each module showed a strongly preserved covariance (Figure 4C), as was the case for the experimental data (Figure 2D). VR adaptation through altered inputs to the motor cortices thus is very similar to the neural activity changes observed *in vivo*.

### Learning through plastic changes within PMd and M1 modules preserves the covariance

Our simulation results so far are consistent with experimental [Tseng et al., 2007, Rabe et al., 2009, Schlerf et al., 2012, Perich et al., 2018] and modelling [Tanaka et al., 2009] studies proposing that VR adaptation is mediated by regions upstream of the motor cortices. But can our model rule out the alternative that adaptation is instead, implemented by recurrent connectivity changes within PMd and M1 (H_local_)?

To test this question, we implemented H_local_ by constraining learning to happen only within PMd and M1, which also led to successful adaptation (Figure 4D). Interestingly, the activity changes produced under H_local_ differed from H_input_ and the experimental data: there were larger changes in the M1 module than in the PMd module (Figure 4E). However, learning based on recurrent weight changes within PMd and M1 did not lead to large changes in covariance, which was largely preserved (Figure 4F), virtually as much as when no local plasticity was allowed (H_input_) (Figure 4C). Thus, the intuition that preserved covariance should be interpreted as a sign of stable underlying connectivity may be misleading.

### Small but coordinated connectivity changes enable motor adaptation

We wished to understand how the model can adapt to the VR perturbation by changing the recurrent connectivity within the PMd and M1 modules without altering their covariance. Interestingly, the connectivity changes under both H_local_ and H_input_ (Figure 5A,B) were small: an average weight change of 1-2% was enough regardless of whether they happened upstream (Figure 5C) or within the motor cortical modules (Figure 5E). These changes were much smaller than those observed during initial training, when the model learned to perform the reaching task from random connection weights (Figure S1B). Thus, “functional connectivity” within the PMd and M1 modules, as measured here by their covariance, may be largely preserved after VR adaptation under H_local_ because network connection weights change very little (Figure S4). We next studied how such small changes in connection weights could nevertheless drive effective behavioural adaptation. Recent studies seeking to relate RNN activity and connectivity have highlighted the importance of low-dimensional structures in connectivity, showing their explanatory power for understanding how tasks are solved [Aljadeff et al., 2016, Mastrogiuseppe and Ostojic, 2018, Schuessler et al., 2020a, Schuessler et al., 2020b]. Inspired by this work, we looked for low-dimensional structures in the connectivity changes emerging in the model during adaptation (Methods). Our analysis revealed that the connectivity change patterns of all plastic modules were low dimensional, independent of where learning happened (Figure 5B,D,F). We hypothesized that the small changes were effective because they were low-dimensional. To test this, we examined how random changes in the connection weights (noise), which are inherently high-dimensional, would affect the behaviour.

**Figure 5:**
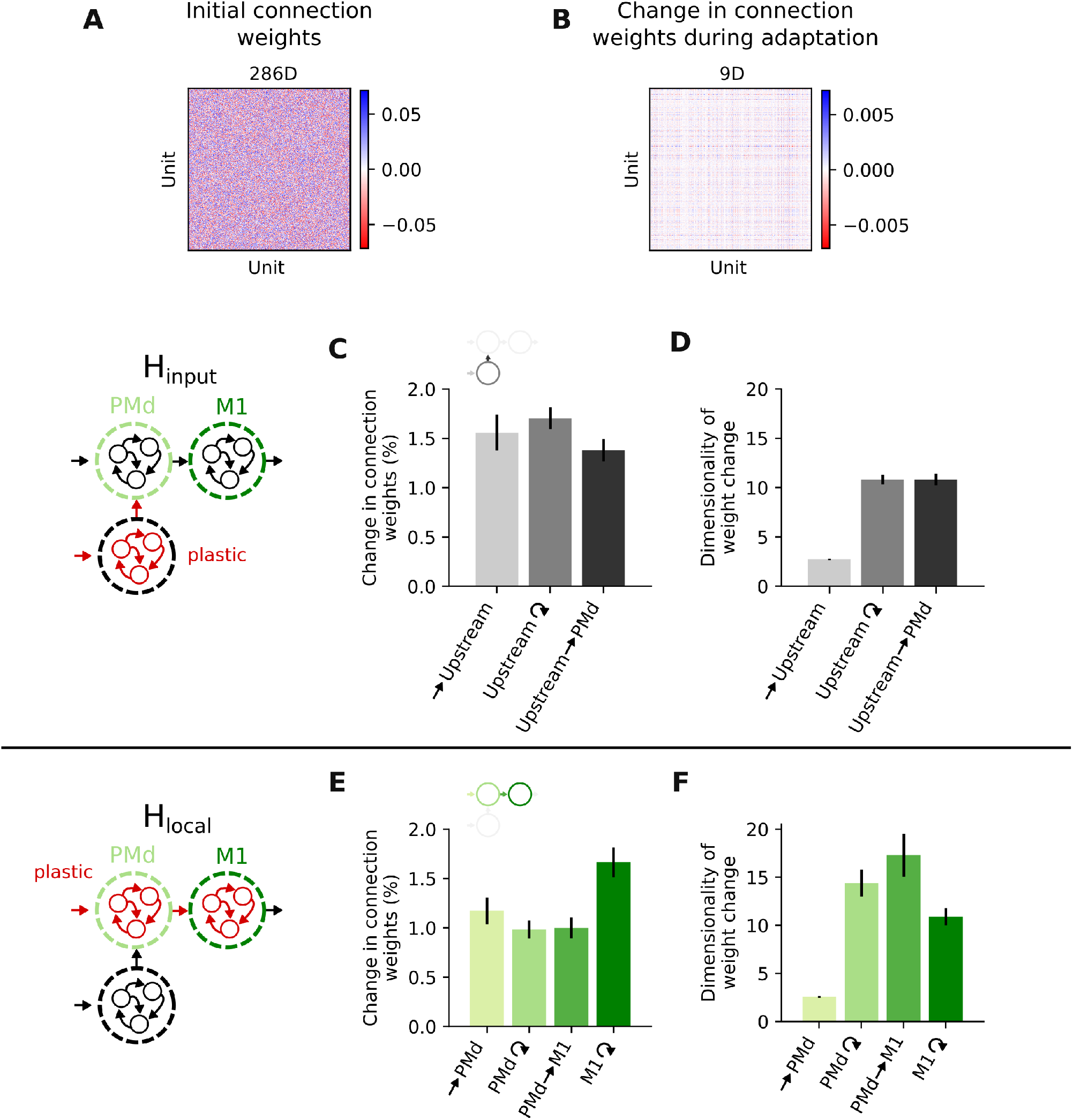
Small but coordinated connectivity changes enable motor adaptation. **A**. Example connection weights for the M1 module after initial training. Top: estimated dimensionality. **B**. Example changes in M1 connection weights following VR learning under H_local_. Same format as A. **C**. Change in connection weights following adaptation under H_input_. Each bar summarizes results for either one module of the network or a set of cross-module connections; bars, mean ± s.d. across 10 network initialisations. **D**. Estimated dimensionality of connection weight changes for each network module and cross-module connections; bars, mean ± s.d. across 10 network initialisations. **E**. Change in connection weights following adaptation under H_local_. **F**. Dimensionality of connection weight changes under H_local_. Data in E,F are presented as in C,D, respectively.

### Low-dimensional connectivity changes are highly robust to noise

For both H_local_ and H_input_, the learned connectivity changes in the model were small and low-dimensional. When considering the biological plausibility of our model, this observation raises the question of how such small connectivity changes could compete with ongoing synaptic fluctuations, which is a known challenge for actual brains [Calvin and Stevens, 1968, Susman et al., 2019, Fauth and van Rossum, 2019, Rule et al., 2020]. To test the hypothesis that the low-dimensionality of the learned connectivity changes is what makes them highly effective, we tested how adding synaptic fluctuations, which are inherently high-dimensional, would affect motor output. Simulating synaptic fluctuations by applying random perturbations to the learned connectivity changes increased the dimensionality of the weight changes (Figure 6B,G), but did not lead to any observable change in reaching kinematics (Figure 6C) or network activity (Figure 6D-E). This was the case even though the applied random perturbations in connectivity were ten times bigger in magnitude than the learned connectivity changes (Figure 6F), completely masking them (Figure 6A-B). Therefore, our model not only suggests that VR adaptation can be implemented based on coordinated synaptic weight changes within PMd and M1, but also that this type of learning would be highly effective due to its robustness to synaptic fluctuations.

**Figure 6:**
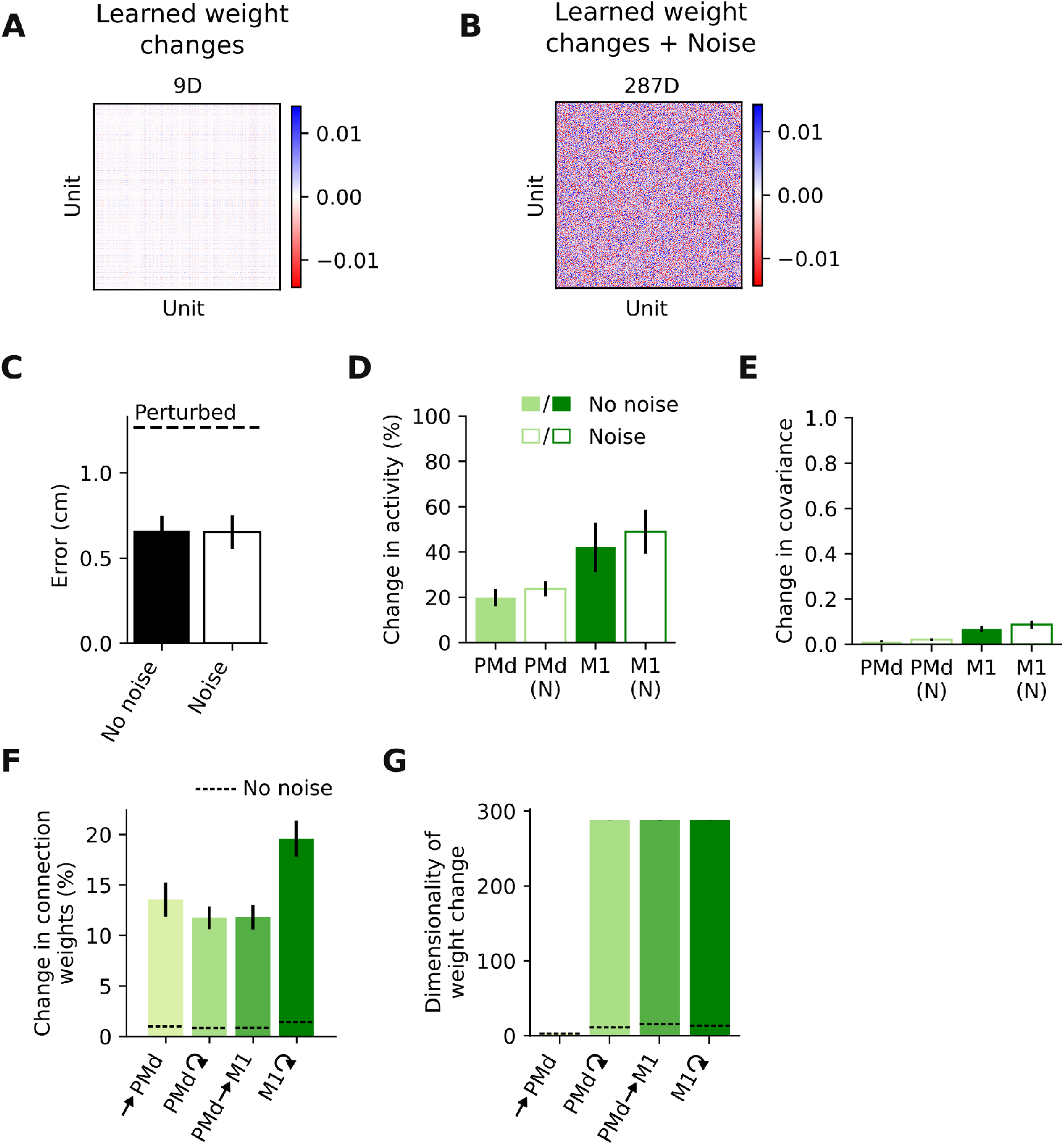
Low-dimensional connectivity changes are highly robust to noise. **A**. Example changes in M1 connection weights following VR adaptation. Same data as in Figure 5B. **B**. Same connection weight changes as in A, but with random connectivity changes added. Note the dramatic increase in the dimensionality of the connection weight changes. **C**. Root mean squared error between target and produced hand trajectories following adaptation in models with and without random weight changes; bars, mean ± s.d. across 10 network initialisations, as in all the panels in this figure. Dashed line shows error under the VR perturbation without any learning. **D**. Change in trial-averaged activity for PMd and M1 module, without (solid) and with (empty) random weight changes. **E**. Change in covariance. Same format as D. **F**. Change in connection weights for each network module and cross-module connections in models with (green bars) and without (dashed lines) noise in connectivity. **G**. Dimensionality of connection weight changes. Same format as F.

### Larger visuomotor rotations allow for a better distinction between H_input_ and H_local_

Although neural activity changes during VR adaptation were better reproduced by a model in which learning happens upstream of the motor cortices (H_input_), activity changes following learning through weight changes within the motor cortices (H_local_) also lined up surprisingly well with the experimental data. To verify that the stable covariance (Figure 4C,F) is not a general feature of the model but reflects task-specific demands, we modelled tasks for which we would expect larger changes.

We first asked the network to learn larger VRs of 60° and 90° instead of the original 30° rotation (Figure 7A). The model was able to compensate for these larger perturbations under both H_input_ and H_local_ (Figure 7B,E). As expected, larger perturbations led to changes in network activity and covariance that increased with rotation angle (Figure 7B,C,F,G). Thus, our model does not necessarily preserve the covariance during learning but adapts according to task requirements. For the 90° rotation, we found a clear difference between H_input_ and H_local_: H_input_ produced larger activity changes in PMd compared to M1, opposite that under H_local_ (Figure 7C,F). Larger rotation angles also increased the difference between H_input_ and H_local_ regarding the covariance changes during learning. Under H_input_, the increase in covariance was similar for the PMd and M1 modules as the rotation increased (Figure 7D). In contrast, under H_local_, the M1 covariance changed more with increasing rotation angle than did that of PMd (Figure 7G). These model predictions could help differentiate between H_input_ and H_local_ in future experiments. In fact, preliminary M1 population recordings obtained during larger VRs (45° and 60°) seemed to match the model predictions for the covariance change under H_input_ (Figure 7D stars), but not H_local_ (Figure 7G stars).

**Figure 7:**
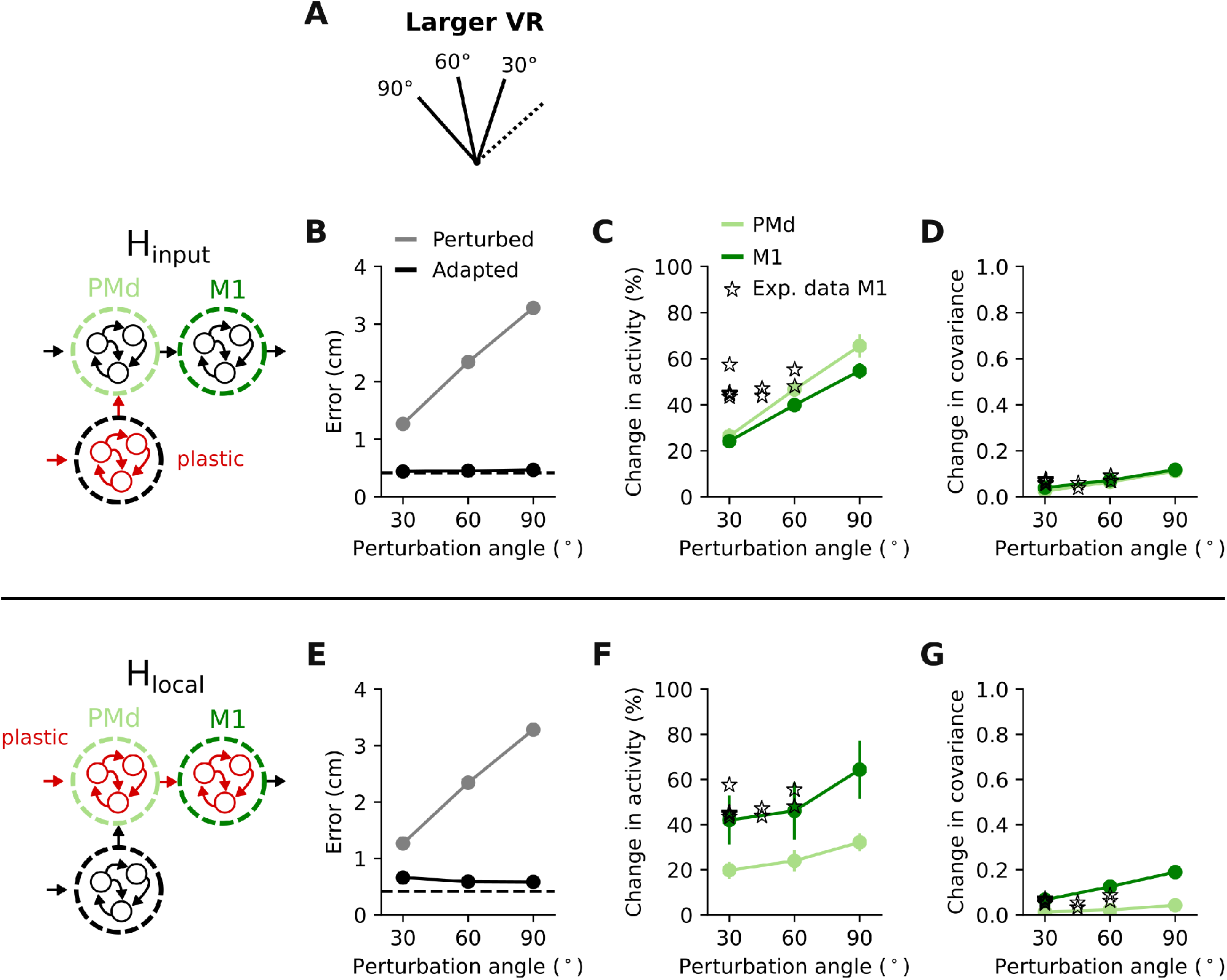
Larger visuomotor rotations allow for a better distinction between H_input_ and H_local_. **A**. To verify that the modelled perturbation does not always produce small activity changes, we tested adaptation to larger VR perturbations (60° and 90°). **B**. Root mean squared error between target and produced hand trajectories without (grey) and with learning (black) under H_input_. Dashed line, error after initial training, with no perturbation applied. **C**. Change in trial-averaged activity for PMd and M1 under H_input_. Markers and error bars, mean ± s.d. across 10 network initialisations, as in all panels in this figure. A few experimental sessions from one monkey with larger rotations are shown as comparison (stars). **D**. Change in covariance following adaptation. Data are presented as in C. Note the similarity between PMd (light green) and M1 (dark green) across all rotation angles. **E**. Error without (grey) and with learning (black) under H_local_. Same format as B. **F**. Change in trial-averaged activity for PMd and M1 under H_local_. **G**. Change in covariance following adaptation. Data in E,F,G are presented as in B,C,D.

### A visuomotor reassociation task can clearly differentiate between H_local_ and learning through reassociation of input signals

Although larger visuomotor rotations help differentiate between upstream learning and learning within the motor cortices, we sought to identify a task that would lead to an even clearer distinction between H_input_ and H_local_. To this end, we implemented a reassociation task where the model had to learn a new, random mapping between cues and reaching directions (Figure 8A; Methods). This task allowed us to test a very specific change in the input signal to the motor cortices that could implement adaptation [Legenstein et al., 2010, Golub et al., 2018]: instead of adjusting the connectivity in an upstream network (H_input_), which allows for highly unconstrained modulation of input signals, the target-related input signals were manually reordered to compensate for the reassociation of cue-reaching direction pairs (Figure 8B). This “learning through input reassociation” resulted in large changes in network activity (Figure 8C), comparable in magnitude to those under H_local_ (Figure 8F). Nevertheless, it did not cause any change in covariance (Figure 8D), which clearly distinguished it from H_local_ (Figure 8G). The reason for this is that, in contrast to H_input_ during VR adaptation, the input signals did not change per se, but were only reassigned to different targets, thereby entirely preserving the network activity patterns.

**Figure 8:**
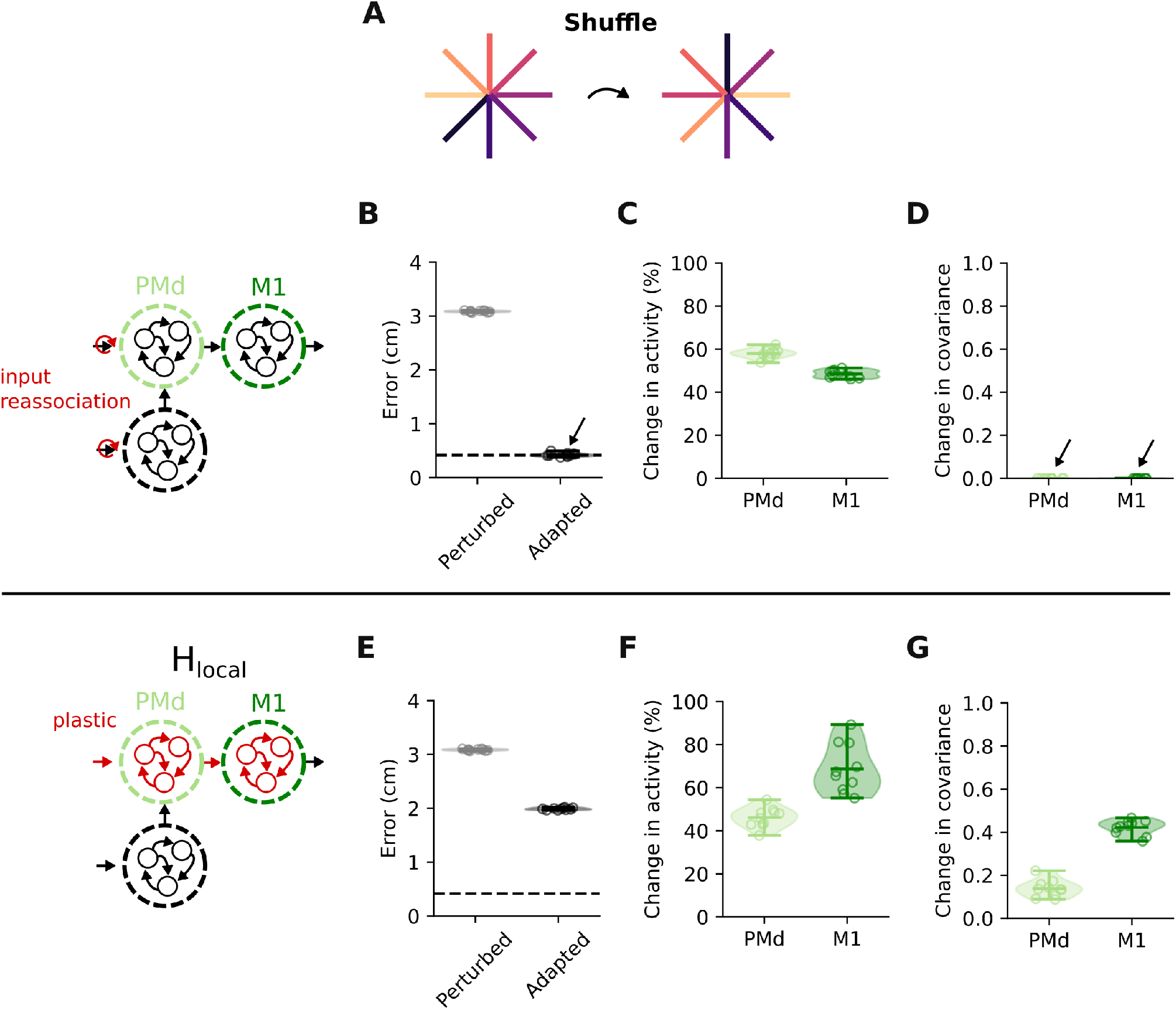
A visuomotor reassociation task can clearly differentiate between H_local_ and learning through reassociation of input signals. **A**. We simulated a reassociation task in which the network had to learn new associations between inputs and reach directions. **B**. Root mean squared error between target and produced hand trajectories without (grey) and with learning (black) through input reassociation. Dashed line indicates error during baseline trials. Markers show 10 different network initializations, shaded area represents estimated distribution, horizontal bars show mean and extrema, as in all panels in this figure. **C**. Change in trial-averaged activity for PMd and M1 under input reassociation. **D**. Change in covariance following adaptation. Note that the covariance matrices do not change at all. **E**. Root mean squared error between target and produced hand trajectories without (grey) and with learning (black) under H_local_. **F**. Change in trial-averaged activity for PMd and M1 under H_local_. **G**. Change in covariance following adaptation. Data in E,F,G are presented as in B,C,D.

## Discussion

Rapid motor learning is associated with neural activity changes in the motor cortices. The origin underlying these changes remains elusive, due to current challenges to measure synapses *in vivo*. Here, we have used modular RNNs simulating the motor cortices to explore whether learning to counteract a VR within tens of minutes could be mediated by local synaptic changes (H_local_). By comparing network activity changes under H_local_ to the changes observed during learning upstream of the motor cortices (H_input_), we have shown how the two hypotheses could be distinguished based on neural population recordings during behaviour. Critically, despite the intuition that learning through plastic changes would lead to detectable changes in neural interactions within and across PMd and M1 populations, both H_local_ and H_input_ (Figure 4) largely preserved the covariance within these two regions, closely matching experimental observations (Figure 2). This likely happened because adaptation under H_local_ was achieved through small, coordinated weight changes within the PMd and M1 network modules (Figure 5). Finally, using our model, we propose tasks for which we anticipate a more dramatic difference between these contrasting hypotheses (Figure 7,8) which can potentially help to interpret experimental data in the future.

Electrophysiological [Tseng et al., 2007, Rabe et al., 2009, Schlerf et al., 2012, Perich et al., 2018] and modelling studies [Tanaka et al., 2009], as well as psychophysical evidence [Thoroughman and Shadmehr, 2000, Criscimagna-Hemminger et al., 2010] suggest that VR adaptation is driven by areas upstream of the motor cortices. Neurophysiological evidence is largely based on the observation that the statistical interactions within PMd and M1 populations remain preserved throughout adaptation [Perich et al., 2018]. This conclusion is in good agreement with studies showing that learning to generate neural activity patterns that preserve the covariance structure only takes a few minutes [Sadtler et al., 2014]. Our direct comparison between H_input_ and H_local_ lends further support to this observation. However, it also paints the intriguing picture that small, globally organized changes in synaptic weights could enable rapid learning without changing the neural covariance, a result that was robust across model initializations (Figure 4) and parameter settings (Figure S2 and Figure S3). Our simulations further indicate that covariance stability is not as directly linked to stable local connectivity as previously thought, as changes in covariance where comparable between H_input_ and H_local_ for a 30° VR perturbation (Figure 4). Instead, the change in neural covariance seemed to be more related to the task itself, as it correlated with the size of the perturbation: the larger the initial error (e.g., caused by larger rotations), the larger the change in covariance (Figure 7). However, the relation between initial error and change in covariance differed depending on where the learning happened (H_input_ or H_local_).

The main difference between the two learning hypotheses we have examined is where in the hierarchical RNN model the connectivity changes occur: within the motor cortices (H_local_), or upstream (H_input_). Although neural covariance was preserved similarly by H_local_ and H_input_, we found a key characteristic that distinguished the two. When local connectivity was allowed to be plastic, the largest activity changes happened within the M1 module, with only small changes in the PMd module (Figure 4E). In contrast, when learning occurred upstream of the PMd and M1 modules, the activity changes were similar in PMd and M1 (Figure 4B). The experimental data, with larger changes in PMd, better matched the pattern produced by H_input_. This observation lends further support for VR adaptation being mediated by plasticity upstream of the motor cortices.

The visuomotor reassociation task allowed us to test a more constrained way in which upstream learning could occur, where input signals were not allowed to change, but were simply reassigned to different targets (Figure 8). Comparing this learning to that mediated by local connectivity changes revealed a clear distinction: learning under H_local_ modified the covariance in both PMd and M1, whereas learning through input reassociation preserved it. Thus, future experiments seeking to disentangle to which extent learning happens within the motor cortices and/or upstream could study this task.

Studies of learning in RNNs have focused on how networks implement *de-novo* training [Hennequin et al., 2014, Sussillo et al., 2015, Rajan et al., 2016, Stroud et al., 2018, DePasquale et al., 2018, Kao, 2019, Yang et al., 2019, Michaels et al., 2020, Schuessler et al., 2020b, Logiaco et al., 2021, Kao et al., 2021]. However, our brain does not learn to perform any task from scratch; it has been “trained” over many generations throughout evolution [Zador, 2019]. Here we studied how neural networks adapt a learned behaviour, as opposed to *de-novo* learning. Our work raises the intriguing possibility that rapid learning following a few tens of minutes of practice could be easily achieved through small but specific changes in circuit connectivity. Thus, initial training seems to provide a highly flexible backbone to adapt behaviour as needed [Goudar et al., 2021].

The fact that the connectivity changes during adaptation under both H_local_ and H_input_ were small and low-dimensional (Figure 5,6) suggests that either one could mediate rapid learning. First, as every synaptic change is costly [Li and Van Rossum, 2020], we would expect a constraint on the total amount of connectivity change in the brain. The VR task being solved with only minor weight changes reflects this; in fact, they could be achieved through long-term potentiation or depression of existing synapses [Froemke and Dan, 2002]. Second, the low dimensionality of these weight changes is also important with respect to solving “credit assignment”, the problem of determining how each synapse should change in order to restore the desired behaviour [Whittington and Bogacz, 2019, Lillicrap et al., 2020, Bellec et al., 2020, Payeur et al., 2021]. Although it is still unclear how this is achieved in the brain, one possibility is that synaptic plasticity is guided by “teacher” signals. Since neuromodulatory signals can regulate synaptic plasticity [Bailey et al., 2000, Nitsche et al., 2006], they seem ideal candidates to regulate biologically plausible learning [Legenstein et al., 2008, Legenstein et al., 2010, Miconi, 2017]. The finding that the connectivity changes needed to adapt to the VR perturbation are “naturally” low-dimensional is promising, as it suggests that learning could be controlled through relatively few neuromodulatory signals. Such implementation would contrast dramatically with the daunting challenge of learning to regulate every single synapse independently.

Lastly, the robustness against synaptic fluctuations conveyed by the low-dimensional connectivity changes makes both H_local_ and H_input_ attractive in terms of ensuring memory stability. Given the fluctuating nature of brain connectivity [Calvin and Stevens, 1968], it remains puzzling how animals remember anything [Susman et al., 2019, Fauth and van Rossum, 2019, Rule et al., 2020, Kossio et al., 2020]. That low-dimensional weight changes, much smaller than ongoing synaptic fluctuations, can achieve successful behavioural adaptation provides a potential solution to this problem.

Our model consistently underestimated the changes in trial-averaged activity observed during VR adaptation, despite closely matching the small covariance changes (Figure 2,4). This is to be expected, as the model only captures changes due to the motor adaptation process itself, whereas the actual neural activity contains signals related to other processes such as impulsivity/engagement [Cowley et al., 2020, Hennig et al., 2021] and feedback processing [Omrani et al., 2016, Stavisky et al., 2017, Kalidindi et al., 2020, Perich et al., 2020, Cross et al., 2021]. In fact, the experimentally observed neural activity changes between the early and late trials of control reaching sessions with no perturbation were almost as large as the changes during adaptation in our model (Figure 2C black dots). How these learning-unrelated changes are combined with the learning-related changes studied here remains unclear. Our modelling predictions for the learning-related changes could help tackle this question in future studies.

Our simulations were not designed to study trial-by-trial learning: we were interested in the neural activity changes between baseline and late adaptation phase when the subjects had largely learned to counteract the perturbation and exhibited stable behaviour (Figure 2B). Given that motor adaptation seems to be mediated by two processes with different timescales [Smith et al., 2006, Huberdeau et al., 2015, Christou et al., 2016], our model mainly captures the slower process of the two. The neural activity changes underlying the early phase adaptation may be driven by different processes [Perich et al., 2018], which our model currently does not test.

In conclusion, our comparison between the activity changes following VR adaptation through motor cortical or upstream plastic changes shows that local plasticity (H_local_) leads to neural signatures that are unexpectedly similar to those of upstream learning (H_input_). Intriguingly, H_local_ not only largely preserved the covariance within PMd and M1 but also resulted in connectivity changes that seem biologically reasonable: they are small, make the network robust against synaptic fluctuations, and can be controlled by relatively few teaching signals. Our simulations thus encourage caution when drawing conclusions from the analysis of neural population recordings during learning, and further suggest potential behavioural tasks that could make it easier to identify where learning is happening.

## Methods

### Tasks

We studied motor adaptation using a visuomotor rotation (VR) paradigm, previously described in Perich et al., 2018 [Perich et al., 2018]. Monkeys performed an instructed delay center-out-reaching task in which they had to reach to one of eight targets uniformly distributed along a circle. All targets were 2 cm squares. The delay period was variable and ranged between 500 and 1,500 ms. For additional details on the task, see [Perich et al., 2018]. During the adaptation phase, visual feedback was rotated clockwise or counterclockwise by 30°, 45°, or 60°. Using our modular RNN model, we simulated both this task and a visuomotor reassociation task in which there was no consistent rotation of the visual feedback; instead, each target required reaching to a different direction, uniquely selected from the initial set of eight different targets.

### Experimental recordings

We analysed eleven sessions from two monkeys (five for Monkey C, six for Monkey M) that were exposed to a clockwise or counterclockwise 30° rotation (data previously presented in [Perich et al., 2018]). In addition to these data, we also analysed three control sessions (one for Monkey C, two for Monkey M) in which no perturbation was applied, as well as additional sessions with larger VR angles from Monkey C where only M1 data was collected (30°, nine sessions; 45°, two sessions; 60°, two sessions) (Figure 7).

The spiking activity of putative single neurons was binned into 10 ms bins and then smoothed using a Gaussian filter (s.d., 50 ms). Only successful trials, where monkeys received a reward at the end, were included in the analysis. We defined the early and late adaptation epochs as the first and last 150 trials of the perturbation phase, when the visuomotor rotation was applied, respectively.

### RNN model

#### Architecture

The neural network contained three recurrent modules, each consisting of 400 neurons, which we refer to as upstream, PMd and M1, respectively (Figure 3A). The PMd and the upstream modules received an identical three-dimensional input signal, with the first two dimensions signalling the x and y target location of that trial, and the third dimension signalling go (1 until the go, and 0 from then on). The upstream module connects to the PMd module and the PMd module connects to the M1 module. The output is calculated as a linear readout of the M1 module activity. Recurrent, as well as feedforward connections were all-to-all. The model dynamics are given by

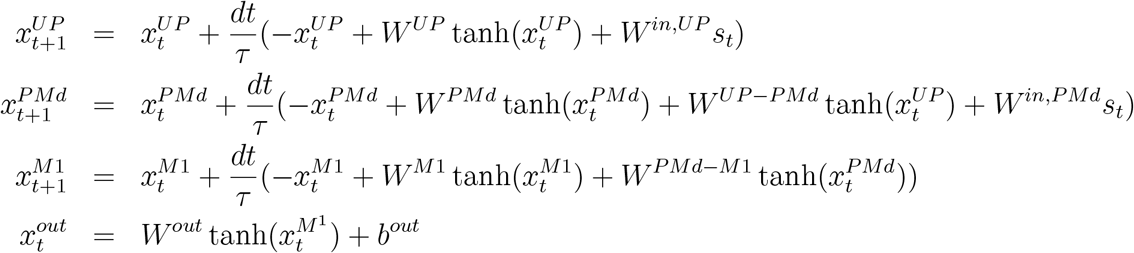

where *x*^*UP*^ describes the network activity in the upstream module, and *x*^*PMd*^ and *x*^*M*1^ the network activity in the PMd and M1 module respectively. *W*^*UP*^, *W*^*PMd*^ and *W*^*M*1^ define the recurrent connectivity matrix within the upstream module, the PMd module and the M1 module, respectively. *W*^*UP*−*PMd*^ defines the connectivity matrix from the upstream module to the PMd module, and *W*^*PMd*−*M*1^ defines the connectivity matrix from the PMd module to the M1 module. The input connectivity matrices for the upstream and the PMd module are given by *W* ^*in,UP*^ and *W* ^*in,PMd*^, respectively; *s*_*t*_ represents the three dimensional input signal described above. The two-dimensional output *x*^*out*^ is decoded from the M1 module activity via the output connectivity matrix *W* ^*out*^ and the bias term *b*^*out*^. The time constant is *τ* = 0.05s and the integration time step is *dt* = 0.01s.

#### Training

Each network was initially trained to produce planar reaching trajectories, mirroring the experimental hand trajectories. The training and testing data set were constructed by pooling the hand trajectories *x*^*target*^ for successful trials during the baseline epochs from all experimental sessions, which resulted in 2238 trials of length 4s (90%/10% randomly split into training/testing). The held out testing data was used to validate that the model had been trained successfully during the initial training period. Model simulations were implemented using PyTorch [Paszke2017] and training was performed using the Adam optimizer [Kingma2014] with a learning rate of 0.0001 (*β*_1_ = 0.9, *β*_2_ = 0.999). The initial training consisted of 500 training trials. The loss function was defined as

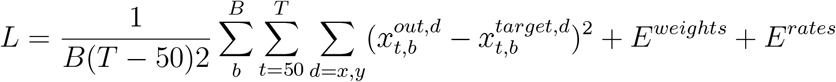

where the regularization term on the weights is given by (∥ ∥ indicates L2 norm)

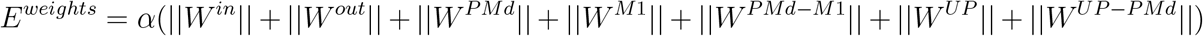

the regularization term on the rates is given by

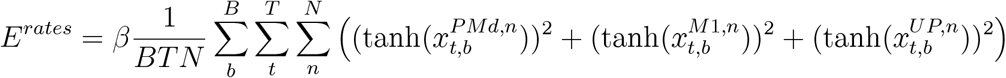

with batch size *B* = 80, time steps *T* = 400 and neurons *N* = 400. The regularization parameters were set to *α* = 0.001, *β* = 0.8. We clipped the gradient norm at 0.2 before we applied the optimization step. For the VR adaptation, we trained the initial network for another 100 trials with the target trajectory rotated 30° (or 60° or 90° for the case of the larger VRs).

### Data analysis

We quantified the changes in actual and simulated neural activity following adaptation using two measures: changes in trial-averaged activity (or peristimulus time histogram, *PSTH*), and changes in covariance. We calculated both metrics within a window that started 600 ms before the go signal and ended 600 ms after it. The change in activity was calculated by

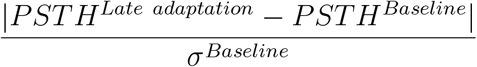

where *PSTH*^*Baseline*^ is the trial-averaged activity in the baseline epoch (experimental data: all baseline trials; simulated data: on a trained model, 100 trials with similar go signal timing), *PSTH*^*Lateadaptation*^ is the trial-averaged activity in the late adaptation epoch (experimental data: last 150 trials of the adaptation epoch; simulation data: on a model trained to counteract the perturbation, 100 trials with similar go signal timing), and *σ*^*Baseline*^ is the neuron-specific standard deviation across time and targets during the baseline epoch. To summarize the change in trial averaged activity across all neurons, time points, and targets, we calculated their median; this provided one single value for each experimental session or simulation run. The change in covariance was calculated using the same trial-averaged data from the baseline and the late adaptation epoch. We calculated the covariance in each of these two epochs and then quantified the similarity by calculating the Pearson correlation coefficient between the corresponding entries of the two matrices. The change in covariance is then defined by 1 minus the correlation coefficient. For the experimental sessions, we computed a lower bound for each measure using the control sessions in which monkeys were not exposed to a perturbation. To account for the fact that there could be activity changes unrelated to motor adaptation [Cowley et al., 2020, Hennig et al., 2021], we compared the activity during 150 consecutive trials from the first half of the control session with 150 consecutive trials from the second half of the control session.

To compute the magnitude of the weight changes after networks learned to counteract the perturbation, we computed the average absolute weight change as

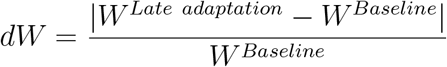

where | | indicates the element wise absolute value, *W*^*Baseline*^ is defined as the model parameter (either *W* ^*in*^, *W*^*UP*^, *W*^*UP*−*PMd*^, *W*^*PMd*^, *W*^*PMd*−*M*1^, or *W*^*M*1^) after the initial training phase but before training on the VR perturbation, and *W* ^*Lateadaptation*^ is defined as the same model parameter after training on the VR perturbation. To obtain one summary value for each simulation run, we calculated the median of all weight entries for a given parameter. To measure dimensionality of weight change we calculated the singular values *s*_*i*_ of *W* ^*Lateadaptation*^ −*W*^*Baseline*^ and defined the dimensionality, using the participation ratio [Gao et al., 2017]:

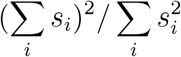

### Statistics

To statistically compare the change in activity found in the control sessions with the change found in the VR sessions, we performed a linear mixed model analysis using R (lmer package). The brain area (PMd or M1) and whether the experimental session included a perturbation phase or not were included as fixed effects, whereas monkey and session identity were included as random effects. A significance threshold of P=0.05 was used.

## Data availability

The data that support the findings in this study are available from the corresponding authors upon reasonable request.

## Code availability

All code to reproduce the main simulation results will be made freely available upon publication on GitHub (https://github.com/babaf/motor-adaptation-local-vs-input.git).

## Acknowledgements

This work has been funded by BBSRC (BB/N013956/1 and BB/N019008/1), Wellcome Trust (200790/Z/16/Z), the Simons Foundation (564408) (all to CC), and the EPSRC (EP/R035806/1 to CC and EP/T020970/1 to JAG). The funders had no role in study design, data collection and analysis, decision to publish, or preparation of the manuscript.

## Supplementary Figures

**Figure S1:**
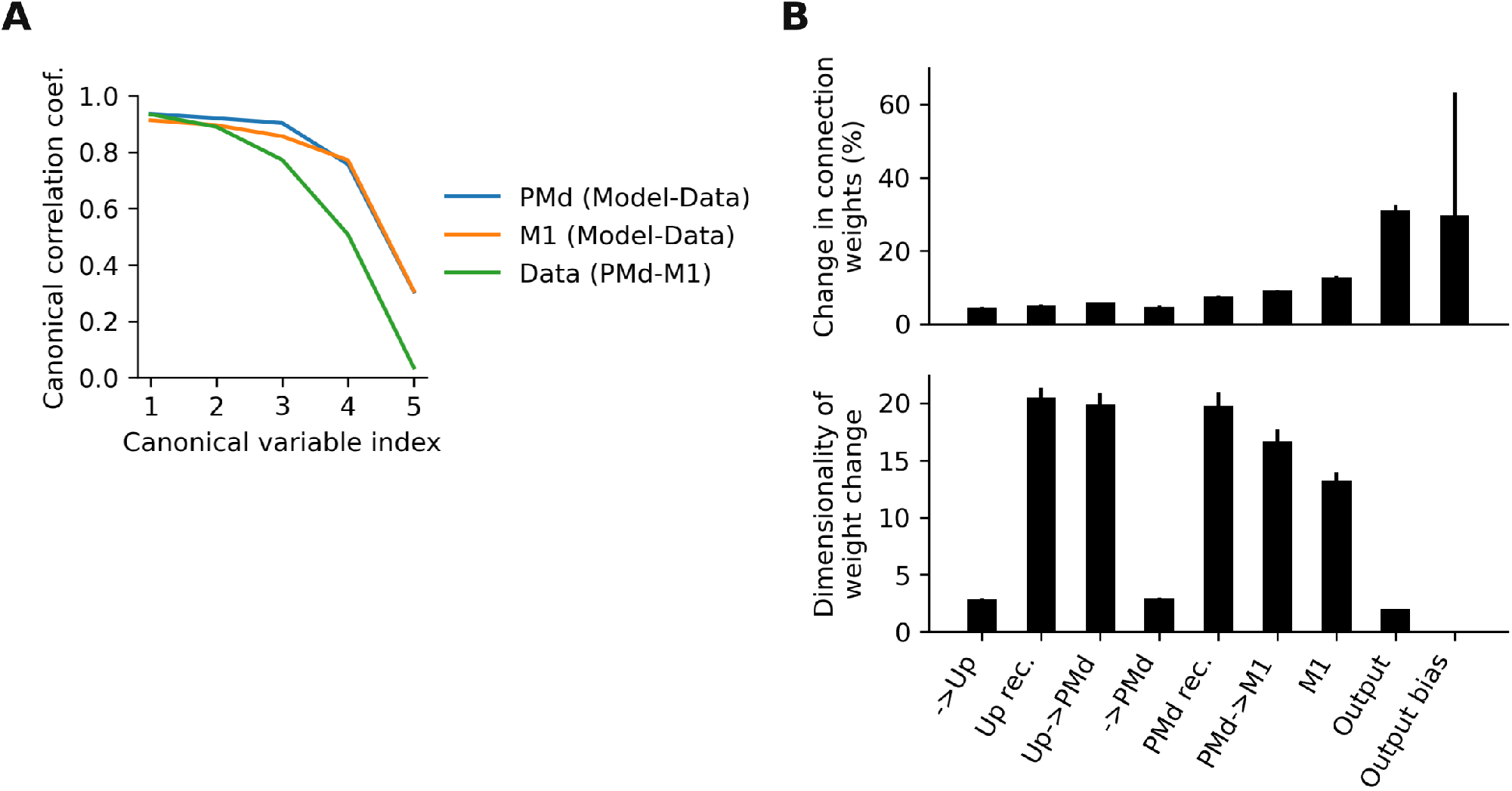
Network activity and weight changes after initial training. **A**. A canonical correlation analysis comparing network activity and neural recordings shows that our model recapitulates key aspects of brain activity: canonical correlations were larger when we compared the corresponding modelled and recorded cortical areas (blue, orange) than when comparing the two experimental brain areas with each other (green). Population activity in the simulated PMd and M1 modules thus largely resemble that of actual neural populations in those areas. Analysis was performed on trial-averaged data which was projected on a 10-D manifold calculated using PCA. **B**. Weight changes (top) and dimensionality of weight changes (bottom) following initial training on the reaching task. Bars and error bars, mean ± s.d. across 10 network initialisations.

**Figure S2:**
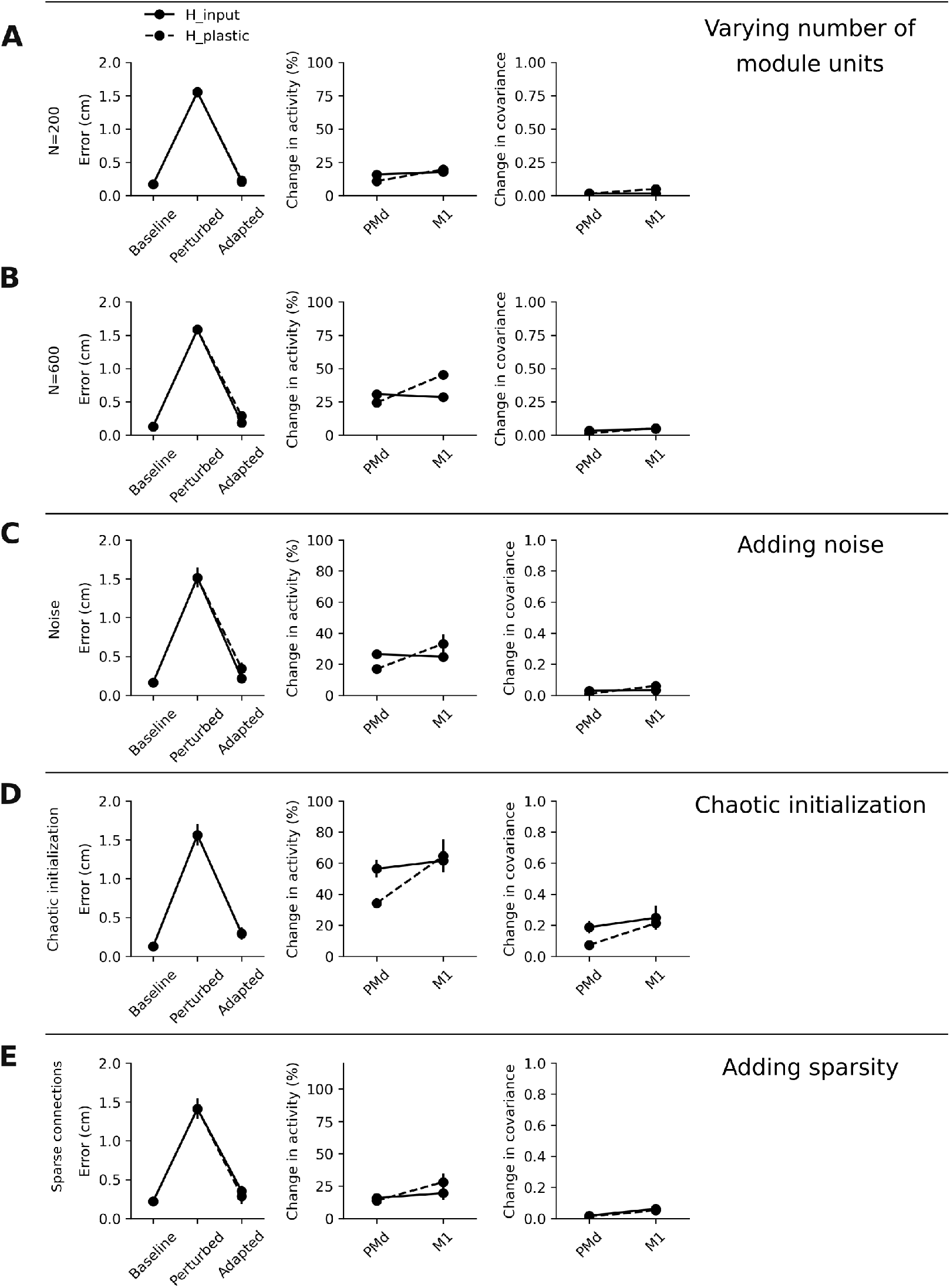
Results are robust to parameter variations. **A**.,**B**. Models with different number of units in each network module exhibit qualitatively similar activity changes. VR adaptation performance, measured as root mean squared error between target and produced hand trajectories (left), change in activity following adaptation (middle), and change in covariance following adaptation (right). Data in B,C,D,E is presented as in A. **C**. Adding noise to the network does not fundamentally change the activity changes in the network. Each neuron in the model received an additional, random, independent input at each time step, drawn from a normal distribution with zero mean and s.d. 0.1. Markers and error bars, mean ± s.d. across 10 network initialisations. **D**. Chaotic initialization slightly increases the overall change in activity and covariance following adaptation, yet preserves the fact that *H*_*input*_ leads to larger changes in PMd compared to M1. Recurrent and inter module weights were initially drawn from a normal distribution with zero mean and s.d. 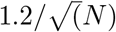. Markers and error bars, mean ± s.d. across 10 network initialisations. **E**. Networks with sparse recurrent and inter module connectivity show similarly low changes in activity and covariance. Only 60% of inter module and 80% of recurrent weights were allowed to be non-zero. Only non-zero connections were plastic during initial training and adaptation. Markers and error bars, mean ± s.d. across three network initialisations.

**Figure S3:**
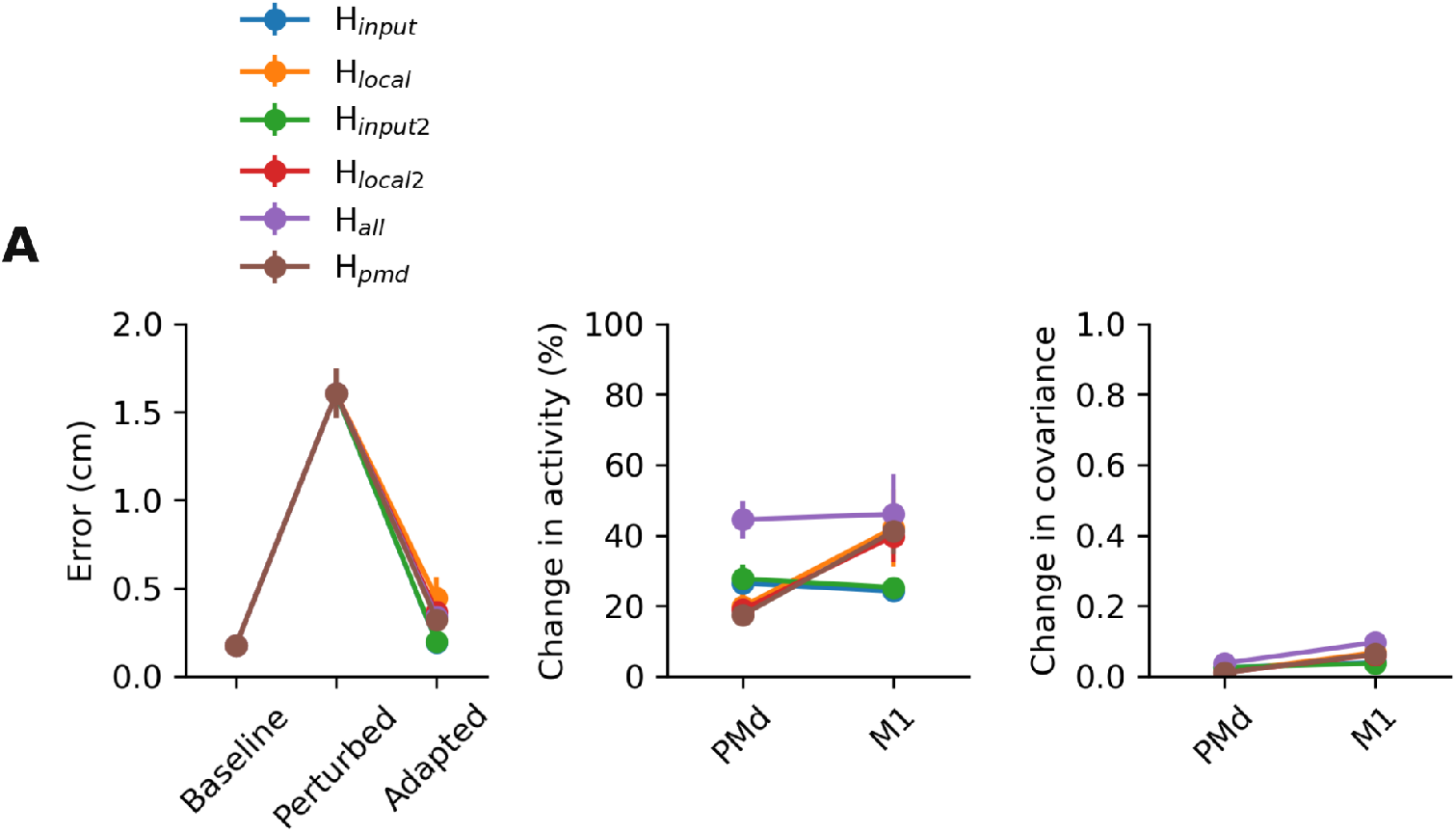
Results are consistent for different plasticity combinations. **A**. VR adaptation performance, measured as root mean squared error between target and produced hand trajectories (left), change in activity between following adaptation (middle), and change in covariance following adaptation (right). Colours indicate which parameters of the model were allowed to be plastic during adaptation. H_input2_ is similar to H_input_ except that the input weight to PMd (*W* ^*in,PMd*^) is also plastic. H_local2_ is similar to H_local_ except that the input weight to PMd (*W* ^*in,PMd*^) is not plastic. For H_all_ every parameter is plastic. For H_pmd_ only the recurrent connectivity within PMd is plastic. Markers and error bars, mean ± s.d. across 10 network initialisations.

**Figure S4:**
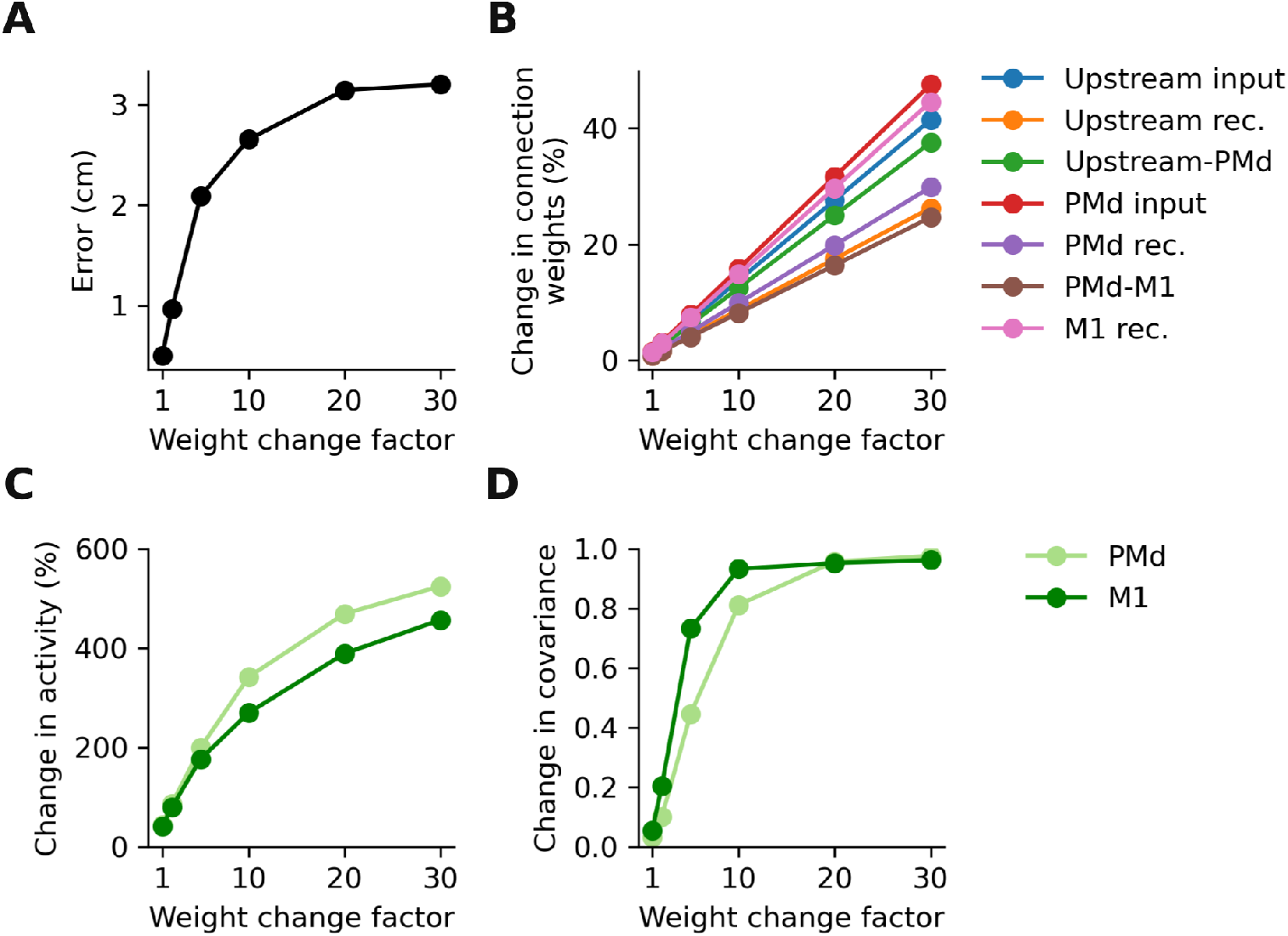
Sensitivity of change in activity and covariance to magnitude of weight change. **A**. VR adaptation performance, measured as root mean squared error between target and produced hand trajectories. Shown is the performance after adaptation for a model where every parameter is allowed to be plastic (H_all_ in Figure S3). To test the sensitivity of the network activity and the produced output on the magnitude of the underlying connectivity changes, we scaled up the learned weight changes during VR adaptation by a factor (x-axis). A weight change factor of 1 thus corresponds to the true weight changes observed after adaptation. Increasing the learned weight changes deteriorates performance, as error increases for increasing weight change factor. **B**. Measured weight change, shown for all model parameters. **C**. Change in activity in the PMd (light) and M1 (dark) modules, respectively. **D**. Change in covariance. Data shown as in C. C,D show that the change in activity and covariance is highly sensitive to the magnitude of the weight change.

## References

Aljadeff, J., Renfrew, D., Vegué, M., and Sharpee, T. O. (2016). Low-dimensional dynamics of structured random networks. Physical Review E, 93(2):022302.

Bailey, C. H., Giustetto, M., Zhu, H., Chen, M., and Kandel, E. R. (2000). A novel function for serotonin-mediated short-term facilitation in aplysia: Conversion of a transient, cell-wide homosynaptic hebbian plasticity into a persistent, protein synthesis-independent synapse-specific enhancement. Proceedings of the National Academy of Sciences, 97(21):11581–11586.

Bellec, G., Scherr, F., Subramoney, A., Hajek, E., Salaj, D., Legenstein, R., and Maass, W. (2020). A solution to the learning dilemma for recurrent networks of spiking neurons. Nature communications, 11(1):1–15.

Calvin, W. H. and Stevens, C. F. (1968). Synaptic noise and other sources of randomness in motoneuron interspike intervals. Journal of neurophysiology, 31(4):574–587.

Christou, A. I., Miall, R. C., McNab, F., and Galea, J. M. (2016). Individual differences in explicit and implicit visuomotor learning and working memory capacity. Scientific reports, 6(1):1–13.

Cowley, B. R., Snyder, A. C., Acar, K., Williamson, R. C., Byron, M. Y., and Smith, M. A. (2020). Slow drift of neural activity as a signature of impulsivity in macaque visual and prefrontal cortex. Neuron, 108(3):551–567.

Criscimagna-Hemminger, S. E., Bastian, A. J., and Shadmehr, R. (2010). Size of error affects cerebellar contributions to motor learning. Journal of neurophysiology, 103(4):2275–2284.

Cross, K. P., Cook, D. J., and Scott, S. H. (2021). Convergence of proprioceptive and visual feedback on neurons in primary motor cortex. bioRxiv.

Das, A. and Fiete, I. R. (2020). Systematic errors in connectivity inferred from activity in strongly recurrent networks. Nature Neuroscience, 23(10):1286–1296.

DePasquale, B., Cueva, C. J., Rajan, K., Escola, G. S., and Abbott, L. (2018). full-force: A target-based method for training recurrent networks. PloS one, 13(2):e0191527.

Diedrichsen, J., Hashambhoy, Y., Rane, T., and Shadmehr, R. (2005). Neural correlates of reach errors. Journal of Neuroscience, 25(43):9919–9931.

Fauth, M. J. and van Rossum, M. C. (2019). Self-organized reactivation maintains and reinforces memories despite synaptic turnover. ELife, 8:e43717.

Feulner, B. and Clopath, C. (2021). Neural manifold under plasticity in a goal driven learning behaviour. PLoS computational biology, 17(2):e1008621.

Froemke, R. C. and Dan, Y. (2002). Spike-timing-dependent synaptic modification induced by natural spike trains. Nature, 416(6879):433–438.

Gao, P., Trautmann, E., Yu, B., Santhanam, G., Ryu, S., Shenoy, K., and Ganguli, S. (2017). A theory of multineuronal dimensionality, dynamics and measurement. BioRxiv, page 214262.

Gerhard, F., Kispersky, T., Gutierrez, G. J., Marder, E., Kramer, M., and Eden, U. (2013). Successful reconstruction of a physiological circuit with known connectivity from spiking activity alone. PLoS computational biology, 9(7):e1003138.

Golub, M. D., Sadtler, P. T., Oby, E. R., Quick, K. M., Ryu, S. I., TylerKabara, E. C., Batista, A. P., Chase, S. M., and Byron, M. Y. (2018). Learning by neural reassociation. Nature neuroscience, 21(4):607–616.

Goudar, V., Peysakhovich, B., Freedman, D. J., Buffalo, E. A., and Wang, X.-J. (2021). Elucidating the neural mechanisms of learning-to-learn. bioRxiv.

Hennequin, G., Vogels, T. P., and Gerstner, W. (2014). Optimal control of transient dynamics in balanced networks supports generation of complex movements. Neuron, 82(6):1394–1406.

Hennig, J. A., Oby, E. R., Golub, M. D., Bahureksa, L. A., Sadtler, P. T., Quick, K. M., Ryu, S. I., Tyler-Kabara, E. C., Batista, A. P., Chase, S. M., et al. (2021). Learning is shaped by abrupt changes in neural engagement. Nature Neuroscience, 24(5):727–736.

Huberdeau, D. M., Krakauer, J. W., and Haith, A. M. (2015). Dualprocess decomposition in human sensorimotor adaptation. Current opinion in neurobiology, 33:71–77.

Kalidindi, H. T., Cross, K. P., Lillicrap, T. P., Omrani, M., Falotico, E., Sabes, P. N., and Scott, S. H. (2020). Rotational dynamics in motor cortex are consistent with a feedback controller. bioRxiv.

Kao, J. C. (2019). Considerations in using recurrent neural networks to probe neural dynamics. Journal of neurophysiology, 122(6):2504–2521.

Kao, T.-C., Sadabadi, M. S., and Hennequin, G. (2021). Optimal anticipatory control as a theory of motor preparation: a thalamo-cortical circuit model. Neuron, 109(9):1567–1581.

Kleim, J. A., Hogg, T. M., VandenBerg, P. M., Cooper, N. R., Bruneau, R., and Remple, M. (2004). Cortical synaptogenesis and motor map reorganization occur during late, but not early, phase of motor skill learning. Journal of Neuroscience, 24(3):628–633.

Kossio, F. Y. K., Goedeke, S., Klos, C., and Memmesheimer, R.-M. (2020). Drifting assemblies for persistent memory. bioRxiv.

Krakauer, J. W., Pine, Z. M., Ghilardi, M.-F., and Ghez, C. (2000). Learning of visuomotor transformations for vectorial planning of reaching trajectories. Journal of Neuroscience, 20(23):8916–8924.

Legenstein, R., Chase, S. M., Schwartz, A. B., and Maass, W. (2010). A reward-modulated hebbian learning rule can explain experimentally observed network reorganization in a brain control task. Journal of Neuroscience, 30(25):8400–8410.

Legenstein, R., Pecevski, D., and Maass, W. (2008). A learning theory for reward-modulated spike-timing-dependent plasticity with application to biofeedback. PLoS computational biology, 4(10):e1000180.

Li, H. L. and Van Rossum, M. C. (2020). Energy efficient synaptic plasticity. Elife, 9:e50804.

Lillicrap, T. P., Santoro, A., Marris, L., Akerman, C. J., and Hinton, G. (2020). Backpropagation and the brain. Nature Reviews Neuroscience, 21(6):335–346.

Logiaco, L., Abbott, L., and Escola, S. (2021). Thalamic control of cortical dynamics in a model of flexible motor sequencing. Cell Reports, 35(9):109090.

Mante, V., Sussillo, D., Shenoy, K. V., and Newsome, W. T. (2013). Context-dependent computation by recurrent dynamics in prefrontal cortex. nature, 503(7474):78–84.

Mastrogiuseppe, F. and Ostojic, S. (2018). Linking connectivity, dynamics, and computations in low-rank recurrent neural networks. Neuron, 99(3):609–623.

Michaels, J. A., Schaffelhofer, S., Agudelo-Toro, A., and Scherberger, H. (2020). A goal-driven modular neural network predicts parietofrontal neural dynamics during grasping. Proceedings of the national academy of sciences, 117(50):32124–32135.

Miconi, T. (2017). Biologically plausible learning in recurrent neural networks reproduces neural dynamics observed during cognitive tasks. Elife, 6:e20899.

Nitsche, M. A., Lampe, C., Antal, A., Liebetanz, D., Lang, N., Tergau, F., and Paulus, W. (2006). Dopaminergic modulation of long-lasting direct current-induced cortical excitability changes in the human motor cortex. European Journal of Neuroscience, 23(6):1651–1657.

Oby, E. R., Golub, M. D., Hennig, J. A., Degenhart, A. D., Tyler-Kabara, E. C., Byron, M. Y., Chase, S. M., and Batista, A. P. (2019). New neural activity patterns emerge with long-term learning. Proceedings of the National Academy of Sciences, 116(30):15210–15215.

Omrani, M., Murnaghan, C. D., Pruszynski, J. A., and Scott, S. H. (2016). Distributed task-specific processing of somatosensory feedback for voluntary motor control. Elife, 5:e13141.

Payeur, A., Guerguiev, J., Zenke, F., Richards, B. A., and Naud, R. (2021). Burst-dependent synaptic plasticity can coordinate learning in hierarchical circuits. Nature neuroscience, pages 1–10.

Paz, R., Boraud, T., Natan, C., Bergman, H., and Vaadia, E. (2003). Preparatory activity in motor cortex reflects learning of local visuomotor skills. Nature neuroscience, 6(8):882–890.

Perich, M. G., Arlt, C., Soares, S., Young, M. E., Mosher, C. P., Minxha, J., Carter, E., Rutishauser, U., Rudebeck, P. H., Harvey, C. D., et al. (2021). Inferring brain-wide interactions using data-constrained recurrent neural network models. bioRxiv, pages 2020–12.

Perich, M. G., Conti, S., Badi, M., Bogaard, A., Barra, B., Wurth, S., Bloch, J., Courtine, G., Micera, S., Capogrosso, M., et al. (2020). Motor cortical dynamics are shaped by multiple distinct subspaces during naturalistic behavior. bioRxiv.

Perich, M. G., Gallego, J. A., and Miller, L. E. (2018). A neural population mechanism for rapid learning. Neuron, 100(4):964–976.

Rabe, K., Livne, O., Gizewski, E. R., Aurich, V., Beck, A., Timmann, D., and Donchin, O. (2009). Adaptation to visuomotor rotation and force field perturbation is correlated to different brain areas in patients with cerebellar degeneration. Journal of neurophysiology, 101(4):1961–1971.

Rajan, K., Harvey, C. D., and Tank, D. W. (2016). Recurrent network models of sequence generation and memory. Neuron, 90(1):128–142.

Rebesco, J. M., Stevenson, I. H., Koerding, K., Solla, S. A., and Miller, L. E. (2010). Rewiring neural interactions by micro-stimulation. Frontiers in systems neuroscience, 4:39.

Rioult-Pedotti, M.-S., Friedman, D., Hess, G., and Donoghue, J. P. (1998). Strengthening of horizontal cortical connections following skill learning. Nature neuroscience, 1(3):230–234.

Roth, R. H., Cudmore, R. H., Tan, H. L., Hong, I., Zhang, Y., and Huganir, R. L. (2020). Cortical synaptic ampa receptor plasticity during motor learning. Neuron, 105(5):895–908.

Rule, M. E., Loback, A. R., Raman, D. V., Driscoll, L. N., Harvey, C. D., and O’Leary, T. (2020). Stable task information from an unstable neural population. Elife, 9:e51121.

Sadtler, P. T., Quick, K. M., Golub, M. D., Chase, S. M., Ryu, S. I., Tyler-Kabara, E. C., Byron, M. Y., and Batista, A. P. (2014). Neural constraints on learning. Nature, 512(7515):423–426.

Schlerf, J. E., Galea, J. M., Bastian, A. J., and Celnik, P. A. (2012). Dynamic modulation of cerebellar excitability for abrupt, but not gradual, visuomotor adaptation. Journal of Neuroscience, 32(34):11610–11617.

Schuessler, F., Dubreuil, A., Mastrogiuseppe, F., Ostojic, S., and Barak, O. (2020a). Dynamics of random recurrent networks with correlated low-rank structure. Physical Review Research, 2(1):013111.

Schuessler, F., Mastrogiuseppe, F., Dubreuil, A., Ostojic, S., and Barak, O. (2020b). The interplay between randomness and structure during learning in rnns. arXiv preprint 2006.11036.

Smith, M. A., Ghazizadeh, A., and Shadmehr, R. (2006). Interacting adaptive processes with different timescales underlie short-term motor learning. PLoS biology, 4(6):e179.

Sohn, H., Meirhaeghe, N., Rajalingham, R., and Jazayeri, M. (2020). A network perspective on sensorimotor learning. Trends in Neurosciences.

Song, H. F., Yang, G. R., and Wang, X.-J. (2017). Reward-based training of recurrent neural networks for cognitive and value-based tasks. Elife, 6:e21492.

Stavisky, S. D., Kao, J. C., Ryu, S. I., and Shenoy, K. V. (2017). Motor cortical visuomotor feedback activity is initially isolated from downstream targets in output-null neural state space dimensions. Neuron, 95(1):195–208.

Stroud, J. P., Porter, M. A., Hennequin, G., and Vogels, T. P. (2018). Motor primitives in space and time via targeted gain modulation in cortical networks. Nature neuroscience, 21(12):1774–1783.

Susman, L., Brenner, N., and Barak, O. (2019). Stable memory with unstable synapses. Nature communications, 10(1):1–9.

Sussillo, D., Churchland, M. M., Kaufman, M. T., and Shenoy, K. V. (2015). A neural network that finds a naturalistic solution for the production of muscle activity. Nature neuroscience, 18(7):1025–1033.

Tanaka, H., Sejnowski, T. J., and Krakauer, J. W. (2009). Adaptation to visuomotor rotation through interaction between posterior parietal and motor cortical areas. Journal of neurophysiology, 102(5):2921–2932.

Thoroughman, K. A. and Shadmehr, R. (2000). Learning of action through adaptive combination of motor primitives. Nature, 407(6805):742–747.

Tseng, Y.-w., Diedrichsen, J., Krakauer, J. W., Shadmehr, R., and Bastian, A. J. (2007). Sensory prediction errors drive cerebellum-dependent adaptation of reaching. Journal of neurophysiology, 98(1):54–62.

Tzvi, E., Koeth, F., Karabanov, A. N., Siebner, H. R., and Krämer, U. M. (2020). Cerebellar–premotor cortex interactions underlying visuomotor adaptation. NeuroImage, 220:117142.

Wang, J., Narain, D., Hosseini, E. A., and Jazayeri, M. (2018). Flexible timing by temporal scaling of cortical responses. Nature neuroscience, 21(1):102–110.

Whittington, J. C. and Bogacz, R. (2019). Theories of error back-propagation in the brain. Trends in cognitive sciences, 23(3):235–250.

Wise, S., Moody, S., Blomstrom, K., and Mitz, A. (1998). Changes in motor cortical activity during visuomotor adaptation. Experimental Brain Research, 121(3):285– 299.

Xu, T., Yu, X., Perlik, A. J., Tobin, W. F., Zweig, J. A., Tennant, K., Jones, T., and Zuo, Y. (2009). Rapid formation and selective stabilization of synapses for enduring motor memories. Nature, 462(7275):915–919.

Yang, G. R., Joglekar, M. R., Song, H. F., Newsome, W. T., and Wang, X.-J. (2019). Task representations in neural networks trained to perform many cognitive tasks. Nature neuroscience, 22(2):297–306.

Zador, A. M. (2019). A critique of pure learning and what artificial neural networks can learn from animal brains. Nature communications, 10(1):1–7.

